# Evidence of opportunistic blood feeding in the parasitic nematode *Heligmosomoides bakeri*

**DOI:** 10.1101/2024.10.09.617485

**Authors:** Edina K. Szabo, James D. Wasmuth, Aralia Leon-Coria, Kaylee D. Rich, Holly Liu, Constance A. M. Finney

## Abstract

Haem, which binds iron and oxygen, is essential for parasitic nematode growth. Nematodes lack endogenous haem synthesis pathways and acquire haem from their environment or intracellular symbionts. Genes involved in haem degradation and detoxification have been identified in parasitic nematodes who are either specialised blood feeders (haematophages) or reside in the blood stream. Targeting these genes, so limiting parasite growth and promoting parasite death, provides a new therapeutic avenue against blood feeding parasitic nematodes.

*Heligmosomoides bakeri*, a model intestinal parasitic nematode, has not been considered a blood feeder. Adults live in the intestinal lumen and graze on host tissue. However, the earlier larval stages enter the intestinal tissue, where they grow and moult multiple times, an oxygen demanding process. We have shown that, *in vivo*, likely through nematode-induced damage, many tissue-dwelling *H. bakeri* are in close vicinity of red blood cells. During this time, infection induces host anemia. Further, tissue-dwelling *H. bakeri* that have fed on blood have an increased expression of collagen genes compared to those parasites that had not fed on blood. *In vitro*, this translates to a growth advantage.

Our findings suggest blood feeding is more widespread among nematodes than currently described. We argue that biochemical adaptations previously considered to be limited to blood-feeders are found in other nematodes. Moreover, *H. bakeri* can serve as a model for studying blood-feeding mechanisms and developing anthelminthic strategies to them.

## INTRODUCTION

Many parasites—including protozoans, helminths, arthropods, and *Cumia* snails—use blood as their primary nutrient source. Among other nutrients, red blood cells are rich in hemoglobin, which when hydrolysed releases large amounts of haem and iron. In blood-feeding (haematophgaus) parasites, haem is an absolute requirement and correlates with parasite growth, survival and reproduction [1]. Nematodes have lost the ability to endogenously synthesise haem and require exogenous/environmental acquisition of haem from their diet, and, in the case of filarial worms, via intracellular symbionts within worm tissues [2]. There are nematodes which are specialised and obligate blood-feeders, such as *Haemonchus contortus* (*Hc*) and the hookworms, including *Ancylostoma caninum* (*Ac*) and *Necator americanus* (*Na*). In the free-living *Caenorhabditis elegans* (*Ce*), haem acquisition—presumably from ingested bacteria—relies on haem-responsive genes (HRGs) [3]. In the lymphatic-dwelling *Brugia malayi* (*Bm*) and gastrointestinal-dwelling *H. contortus*, the HRGs so far identified and verified are more highly expressed in larval stages than adult [4,5]. Since the primarily blood-feeding stage of *H. contortus* is the adult rather than the larval stage, it is likely that other *Hc*-HRGs await discovery.

Nematodes also lack the genes coding for haem oxygenase-based haem degradation, a common method used by other blood-feeding parasites to detoxify haem and release iron [2]. However, the ability of specialist blood feeding nematodes to degrade hemoglobin and detoxify haem has been intensely studied [6–8]. The aspartic proteases, APR-1 and APR-2 (*Ac*-APR-1 from the dog hookworm *Ancylostoma caninum*, *Na*-APR-1 and *Na*-APR-2 from the human hookworm, *N*. *americanus)*, can digest hemoglobin *in vitro* [9]. The glutathione S-transferases, *Ac*-GST-1 and *Hc*-GST1 contain a high-affinity binding site for hematin and haem-related compounds [7,8]. Both APR-1 and GST-1 are the focus of anti-hookworm vaccine research and the subject of clinical trials [10,11]. Other genes of interest are *Ac*-CP-2 that also digests hemoglobin [30] and *Ac*-MEP-1 that acts downstream of *Ac*-CP-2 and *Ac*-APR-1 [31,32]. They have been successfully trialed as vaccine targets in dogs [33] and hamsters [34] respectively. Interestingly, research demonstrating that the free-living *C. elegans* can feed on blood *in vitro* with no detrimental effects [12] suggests that the ability to blood feed may be more widespread than initially thought and access to blood *in vivo* may be a key factor in determining whether a nematode uses blood as a regular food source.

*Nippostrongylus brasiliensis* (*Nb*) is a model of human hookworm infection due to its similar lifecycle—migration through the lungs prior to adult residence in the small intestine—and its ability to ingest blood [13]. However, *N. brasiliensis* is phylogenetically removed from the hookworms and its blood-feeding is likely limited to larval and immature adult stages (Fig 1A) [14]. The mouth morphology of adult *N. brasiliensis* suggest that they do not acquire host blood (compare with the stylet of *H. contortus* or the cutting plates of *N. americanus*) [15]. Also, in the mouse, *N. brasiliensis* adults are not able to remain in the host for more than ∼10-14 days, making it a relatively short infection compared to the chronic human hookworm infections. *Heligmosomoides bakeri* (*Hb*) is a close relative to *N. brasiliensis* (Fig 1A) and a widely used rodent gastrointestinal model parasite. For many years, *H. bakeri* and *Heligmosomoides polygyrus* were mistakenly regarded as the same species. However, genomic comparisons and differences in host specificity have since confirmed that they are distinct species [16]. Similar to *H. contortus*, the *H. bakeri* lifecycle is entirely enteric (Fig 1B). Infective stage 3 larvae are ingested by their host and exsheath in the stomach. They migrate to the small intestine and enter the intestinal mucosa, in which they develop over the course of ∼8 days. They exit as adults into the lumen and reside chronically coiled around the intestinal villi but do not cause appreciable damage to tissue [17]. Previous work identified the food source of adult *H. bakeri* as host tissue, ruling out blood and food ingesta [18]. However, feeding of the *H. bakeri* tissue dwelling stages—stage 3 and 4 larvae, and newly moulted immature adults—has not been characterised. With two moults and considerable growth (parasites can triple in size), understanding the food source of tissue dwelling stages is key to understanding the parasite’s needs in this niche.

**Figure 1:**
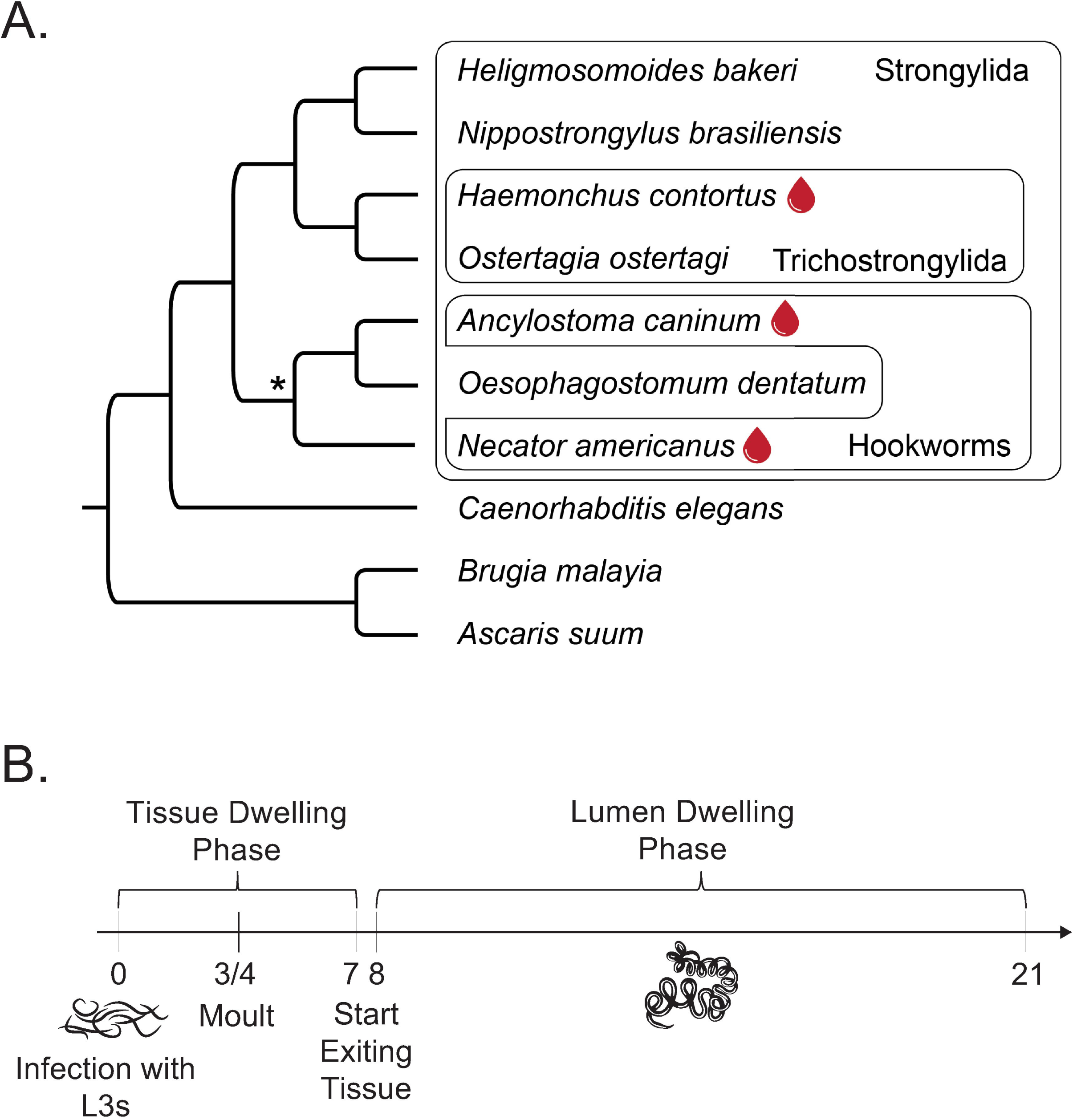
The gastrointestinal nematode *H. bakeri.* (A) Phylogenetic relationship of species described in this study. The widely accepted specialised blood feeders are denoted by the blood drop. The tree’s topology is the widely accepted nematode species tree, as determined by many studies. One area of disagreement is the monophyly of the hookworms, denoted by *. We present the topology that has most frequently reported, but acknowledge that other studies report a hookworm monophyly. (B) *H. bakeri* infection timeline. C57Bl/6 mice are infected orally with infective L3s on D0. The larvae exsheath and enter the small intestinal tissue within a day. At 3 to 4 days post-infection, parasites moult in the intestinal tissue. At day 7/8 post-infection, parasites start exiting the tissue and living in the lumen coiled around the intestinal villi. Adults remain in the lumen for at least 21 days.

Using feeding and growth assays, as well as transcriptomic profiling, we have demonstrated that *H. bakeri* larvae and adults can feed on red blood cells *in vitro*, and that this blood meal provides a growth advantage to developing larvae but not to adults. *In vivo,* tissue-dwelling nematodes—larvae and newly moulted adults—have access to blood, and we show that ingesting blood is associated in increased gene expression of cuticle forming collagen and oxygen carrying globin genes as well as *APR-1*, *GST-1* and *HRG*s. However, once they exit tissue into the lumen, the growth advantage afforded by blood feeding is no longer apparent. These results demonstrate that facultative haematophagy is potentially common in nematodes, and *H. bakeri*, which is easier to maintain *in vitro* and *in vivo* than specialised blood feeders, could be used during the initial stages of anthelminthic development targeting blood feeding.

## METHODS

### Mice and parasites

C57Bl/6 mice, 6-8 weeks of age were used for all experiments, bred and maintained in the Life and Environmental Science Animal Resource Centre (LESARC) in the Department of Biological Sciences, University of Calgary, Canada. Original stocks were obtained from Charles River, Canada. Mice were acclimatised for a minimum of two days when moved to our experimental room and housed in groups of 3-5 in static cages. All animal experiments were approved by the University of Calgary’s Life and Environmental Sciences Animal Care Committee (protocol AC18-0168). All protocols for animal use and euthanasia were in accordance with the Canadian Council for Animal Care (Canada). Enrichment was provided in the form of plastic housing, bedding.

Infected mice were orally gavaged with 200 or 400 third stage *Heligmosomoides bakeri* larvae (original stock gifted by Dr. Allen Shostak, University of Alberta, Canada, further experiments conducted with a stock from Dr Lisa Reynolds, University of Victoria) and euthanized at different time points post infection. The *H. bakeri* lifecycle was maintained according to published methods [19]. Larvae were obtained from fecal cultures after approximately 8-9 days of incubation at room temperature.

L3 larvae were exsheathed using 0.02% NaOCl at 37 °C for 80 min to obtain activated exsheathed larvae. Adult worms were removed from the intestinal tract and isolated using a modified Baerman apparatus. Exsheathed larvae and adults were washed three times with water and placed in Dulbecco’s modified eagle’s medium – high glucose (Sigma cat. D5796) where they were counted; adults were also sexed. Worms were either used immediately for feeding assays or snap frozen and kept at -80°C until RNA isolation.

Photographs were taken at various post-infection time points using a light microscope (DMRB Leica)/Qimaging micropublisher 3.3 rtv and a Leica DMRB Microscope and fluorescent microscope/camera Zeiss AxioVert 200M Fluorescence Microscope.

### Tissue Histology

Consecutive formalin fixed paraffin embedded mouse small intestinal sections were stained with H&E by the University of Calgary’s Veterinary Diagnostic Services Unit. Photographs of worms were taken at x400 magnification using a light microscope (DMRB Leica). Red blood cells were identified by their unique staining patterns.

### In vivo hematocrit assessment

Hematocrit levels of mice were measured at different time points post-infection. Blood was collected from each mouse’s tail vein directly into heparinized glass hematocrit capillary tubes. Tubes were centrifuged for 15 min at 1500xg. The packed cell volume of each capillary tube was measured with a ruler to the nearest millimeter. Hematocrit levels were calculated as a percentage of packed red blood cells to the total length occupied by the whole blood. The hematocrit for each animal was measured at baseline (D0). Later measurements were matched back to this baseline value.

### In vitro feeding assays

Fluorescent Beads: Exsheathed larvae and adult worms were cultured in the presence/absence of 0.5 mm FITC labelled and/or 6 mm PE labelled fluorescent beads (Polysciences) for 5 days at 0.5µM (1 x10^7^ beads/ml). Photographs of the exsheathed larvae/adult worms were taken using a Zeiss AxioVert 200M fluorescent microscope. The number of live exsheathed larvae/adult worms having ingested beads was counted as a percentage of total live worms in the well and the fluorescent intensity of the worms was calculated using ImageJ.

Red Blood Cells: Mouse blood was collected with Heparin (Sandoz, 10750). 20 drops of Alsever’s solution were added to inhibit clotting. The blood was washed and suspended in a balanced salt solution: KCl (0.035%), CaCl2 (0.029%), MgSO4 (.029%), NaCl (0.82%), Tris-base (0.25%), glucose (0.1%), BSA (0.5%), pH 7·4. Red blood cells were counted using a haemocytometer, and used at a final concentration of 2 x 10^8^ cells per ml. Both exsheathed larvae and adult worms were incubated for 2 or 5 days. At the end of the experiment, exsheathed larvae were observed under a light microscope (DMRB Leica) to assess the presence/absence of pigmentation. Adult worms were pelleted and resuspended in NaOH for 24 hours to allow the worms to dissolve. Absorbance of the resulting solution was measured using a VMax Kinetic ELISA Microplate Reader with Softmax Pro Software at a wavelength of 400 nm.

Quinidine powder (Sigma, Q3625) was resuspended in DMSO. Worms were cultured in the presence/absence of quinidine (100 μM).

### RNA sequencing

Methods for obtaining the data are published [20]. Briefly, RNA was isolated using Trizol and the Miniprep kit (Zymo, R2050) following the manufacturer’s instructions. The RNA was then cleaned using the Zymo RNA Clean and Concentrator -5 kit (Zymo, R1015) following the manufacturer’s instructions. Quantity and quality of the total RNA was assessed on an Agilent 2200 TapeStation RNA ScreenTape following the manufacturer’s instructions. Libraries were made from suitable RNA samples using the NEBNext Ultra II Directional RNA Library Prep Kit following the manufacturer’s instructions. Libraries were multiplexed and paired-end sequenced on an Illumina NovaSeq with a S2 flow cell or SP flow cell for 300 cycles (2×150bp) using the v1.5 reagent kit, following the manufacturer’s instructions. The resulting reads have been deposited in the SRA and are awaiting an accession number. RNA-seq reads were aligned to the *H. bakeri* genome assembly (PRJEB15396) obtained from WormBase ParaSite release 17 (WBPS17) using STAR v.2.7.11b [21]. Read counts per transcript were calculated using featureCounts v.2.0.6 [22] and used for differential expression analysis with the R package DESeq2 v.1.42.0 [23]. Linux commands and R code can be found at https://github.com/Kayleerich/hbakeri_bloodfeeding.

### Gene and protein identifiers

The following genes and proteins are described in this manuscript. In Table 1, we provide their database identifiers and orthologue information to assist with study reproducibility.

**Table 1:** Information on genes described in this study.

| Gene | Species | Uniprot ID | Genbank ID | Wormbase ID<sup>†</sup> | Ortholog information<sup>†</sup> |
| --- | --- | --- | --- | --- | --- |
| *Bm-hrg-1* | *Brugia malayi* | A0A0K0JHU8 | XP_042931527 | Bm5182 | *Ce*: WBGene00009493, WBGene00009494, WBGene00009495, WBGene00019830; *Hb*: HPOL_0001917901; *Nb*: Nippo_chrX.g15524; *Hc*: HCON_00169910 |
| *Bm-hrg-2* | *Brugia malayi* | A0A8L7SYH0 | XP_042933895 | Bm2383 | *Ce*: WBGene00000412, WBGene00008296, WBGene00008297, WBGene00010470, WBGene00010471, WBGene00010472, WBGene00010473, WBGene00016116; *Hb*: HPOL_0000121101, HPOL_0000497501, HPOL_0002148701; *Nb*: Nippo_chrII.g4342, Nippo_chrIII.g6108, Nippo_chrIII.g6109, Nippo_chrIII.g6113; *Hc*: HCON_00074830, HCON_00074850, HCON_00074860, HCON_00074865, HCON_00074867, HCON_00074870, HCON_00074880, HCON_00074900, HCON_00192970 |
| *Hc-hrg-2* | *Haemonchus contortus* | A0A7I5E8W1 | MK371241 | HCON_00074880 | See orthologue information for *Bm-hrg-2* |
| *Na-apr-1* | *Necator americanus* | Q9N9H3 | AJ245459 | Necator_chrX.pre1.g25887 | *Ce*: WBGene00000217; *Hb*: HPOL_0001873801; *Nb*: Nippo_chrX.g15713; *Hc*: HCON_00103350 |
| *Na-apr-2* | *Necator americanus* | Q9N9H4 | AJ245458 | Necator_chrIV.pre1.g17227 | *Ce*: none; *Hb*: HPOL_0000103201; *Nb*: Nippo_chrV.g14115, Nippo_chrV.g14122; *Hc*: HCON_00153270, HCON_00153310 |
| *Ac-gst-1* | *Ancylostoma caninum* | A0A368H0I6 | AY605283 | ANCCAN_04558* | *Ce*: WBGene00001754; *Hb*: HPOL_0001908401, HPOL_0001908501, HPOL_0001908601, HPOL_0002218401; *Nb*: Nippo_chrII.g4085, Nippo_chrII.g4086, Nippo_chrII.g4087, Nippo_chrIV.g11275 |
| *Ac-mep-1* | *Ancylostoma caninum* | A0A368FLE5 | AF273084 | ANCCAN_21297* | *Ce*: WBGene00020788; *Hb*: HPOL_0001315501; *Nb*: none |
| *Ac-cp-1* | *Ancylostoma caninum* | A0A368GSJ8 | U18912 | ANCCAN_06644 | *Ce*: none ; *Hb*: none; *Nb*: none; *Hc*: none |
| *Ce-glb-1* | *Caenorhabditis elegans* | P30627 | 176261 | WBGene00014030 | *Hb*: 14 genes; *Nb*: 12 genes; *Hc*: 13 genes |

<sup>†</sup> These identifiers were taken from either Wormbase (https://wormbase.org/) or Wormbase Parasite (https://parasite.wormbase.org)
‡ The ortholog information was taken from Wormbase Parasite (https://parasite.wormbase.org/). In several cases, the orthologs were in a 1-to-many or many-to-many relationship. We report all genes considered orthologs / co-orthologs. For specific details, see the relevant gene page on Wormbase Parasite
\* In some instances, the gene model presented on Wormbase Parasite is noticeably shorter than the entry in Genbank.

### Statistical analysis

Linearity was assessed using a normality test when the n was sufficiently high. For nonparametric data, Mann Whitney/Kruskal Wallis tests with Dunn’s multiple comparisons were used to assess differences between either two or more experimental groups using GraphPad Prism. For parametric data, we used T-tests/ANOVA with Sidak’s multiple comparisons. Two-way ANOVAs were used when comparing experimental means and their standard deviations. On graphs, the line represents the median unless otherwise stated.

## RESULTS

### *In vitro, H. bakeri* larvae and adults can ingest blood

When cultured in blood, most exsheathed larvae had visibly darker digestive tracks (94%, SD +/- 3%, Figs 2A and 2B), a hallmark of RBC ingestion [13]. As adult *H. bakeri* are naturally pink in colour, changes in pigmentation due to RBC ingestion are harder to observe reliably by eye.

**Figure 2:**
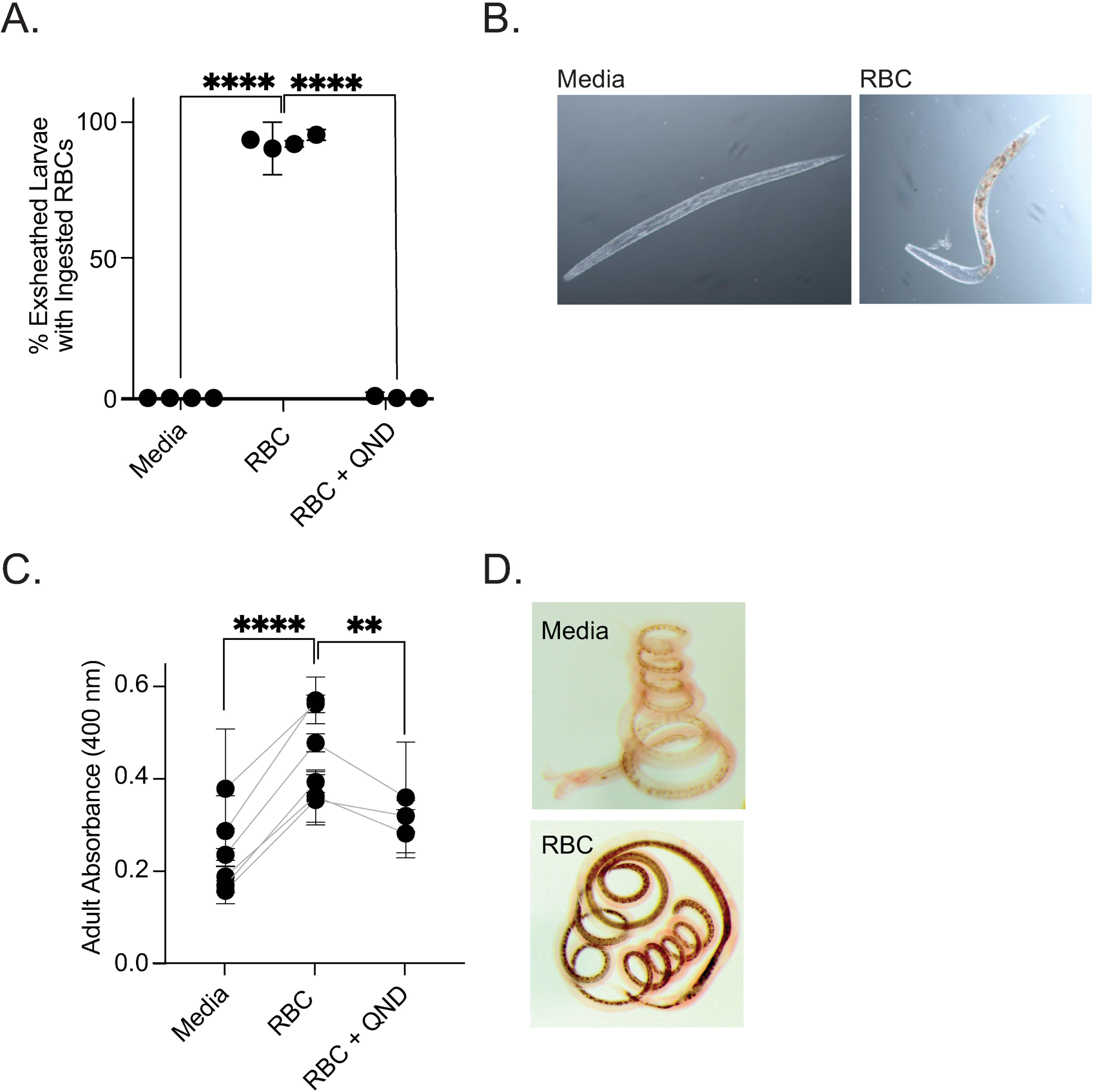
*H. bakeri* ingesting blood *in vitro.* (A-B) Exsheathed larvae were cultured *in vitro* in triplicate with media or 2 x 10^8^ cells per ml murine red blood cells (RBC) for 5 days in the presence or absence of 100 mM QND. (A) The percentage of pigmented larvae. Each dot represents one experiment which is the mean value of 3 replicate wells containing a minimum of 25 and a maximum of 147 worms. (B) Photographs of representative exsheathed larvae after culture. (C-D) C57Bl/6 mice were infected orally with 400 infective L3s. At D10, worms were removed from the intestinal lumen and 20 adults were cultured *in vitro* in triplicate per experiment. Adults were cultured with media or 2 x 10^8^ cells per ml murine red blood cells (RBC) in the presence or absence of 100 mM QND. (C) After 5 days of incubation, worms were collected and incubated in 0.1 M NaOH for 24hrs. Absorbance was measured at 400 nm. Each dot represents one experiment which is the mean value of 3 replicate wells containing 20 worms. (**D**) Representative photographs of adults after culture. Two-way ANOVA, n.s. = non significant, * = p<0.05, **** = p<0.0001.

Therefore, as well as observing a shift in colour, we also measured their absorbance, which were both consistent with adults ingesting RBCs (Figs 2C and 2D). It has been suggested that quinidine, a Na^+^ channel blocker [24], affects a specific blood feeding-related pathway [13]. In the presence of quinidine, RBC ingestion stopped in *H. bakeri* larvae and reduced significantly in adults (Figs 2A and 2C) supporting similar findings with *N. brasiliensis* [13]. However, using a bead feeding assay, we show that ingestion of beads in *H. bakeri* larvae and adults was also reduced in the presence of quinidine (Sup Figs 1 and 2), confirming quinidine’s previously described broader effects on nematodes’ ability to feed [24].

### *In vivo, H. bakeri* tissue-dwelling stages induce anemia

We show that on days 4-to-6 post-infection (D4-6, tissue-dwelling phase), *H. bakeri* are in close contact with blood which pools around the worm, likely because of worm-induced damage (Fig 3A and 3B). We attempted to view RBC ingestion *in vivo* by injecting mice with fluorescently labelled RBCs, but could not detect fluorescent worms. This was probably due to the small number of RBCs ingested over a long period of time (∼7 days) along the entire small intestine. However, we did observe a small but significant anemia—measured by a reduction in hematocrit at D7—when *H. bakeri* are tissue-dwelling (Fig 2D). The anemia was no longer apparent when worms had left the tissue and were in the lumen (Fig 2E).

**Figure 3:**
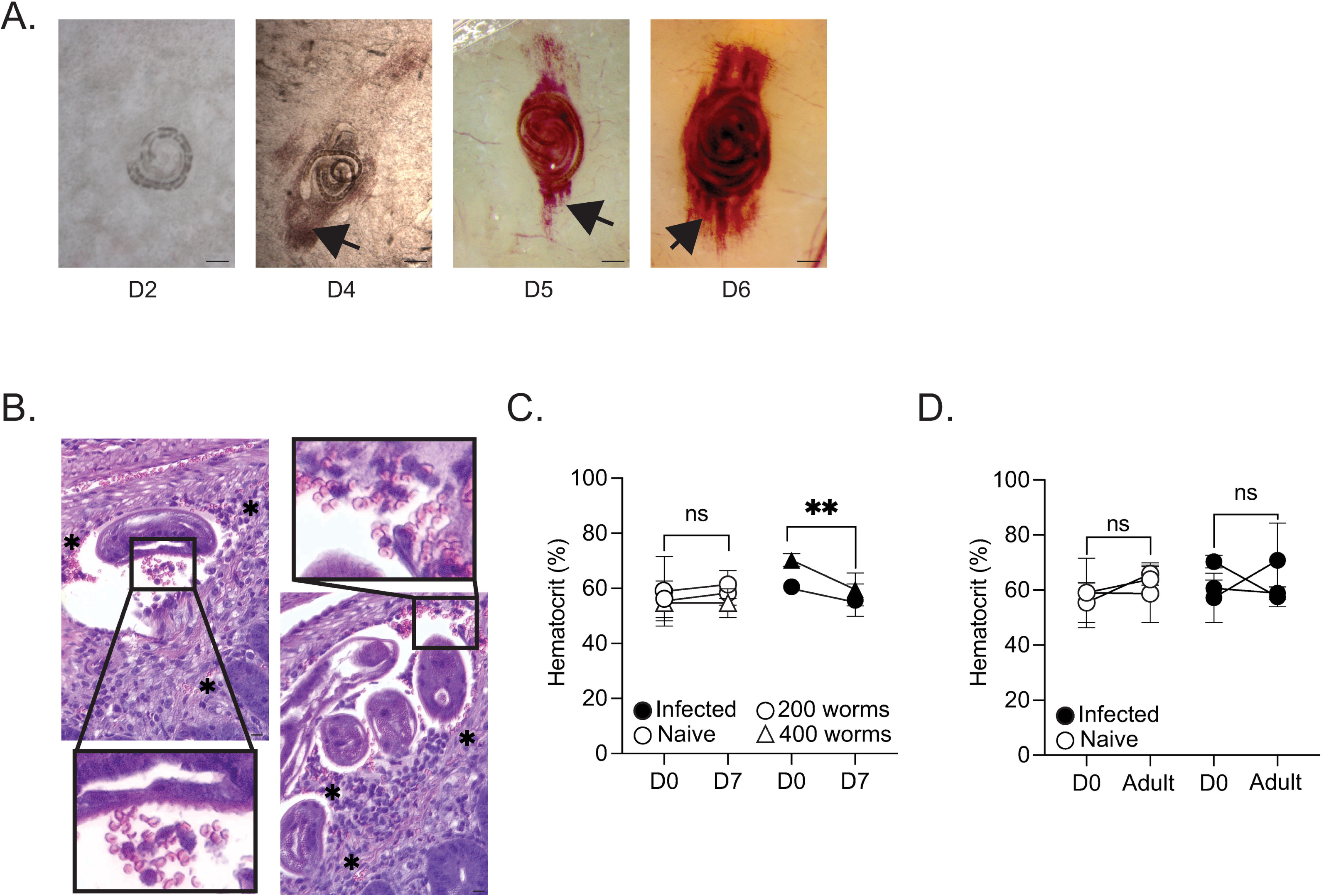
Tissue stage *H. bakeri* blood feeds *in vivo.* C57BL/6 mice were infected with 400 *H. bakeri* larvae. Representative photographs of *H. bakeri* tissue dwelling stages at 2, 4, 5 and 6 days post-infection. Black arrows depict accumulations of mouse red blood cells around the parasite. Scale = 0.1mm for days 2 and 4, 1mm for days 5, 6 and 7. (C) C57BL/6 mice were infected with 200 *H. bakeri* larvae. Representative photographs of *H. bakeri* tissue dwelling D4 stages. Black boxes show close up of red blood cells (in pink). Symbols * depict other accumulations of mouse red blood cells around the parasite. (D) C57BL/6 mice were infected with either 200 (circles) or 400 (triangles) *H. bakeri* larvae and hematocrit measured for each animal at D0, D7 and D11-12 (adult) post-infection in naïve and infected mice. Each dot represents one experiment which is the mean value of a minimum of 5 and a maximum of 16 mice.

### Blood feeding provides a growth advantage for tissue dwelling stages *in vitro*

To assess whether RBC ingestion provides a fitness advantage to the exsheathed larvae, they were measured after a two-day incubation. Larvae having ingested RBCs (D2 RBC), increased in length compared to those cultured in media alone or in the presence of RBCs and QND (D2 Media, D2 RBC + QND, Fig 4A and 4B).

**Figure 4:**
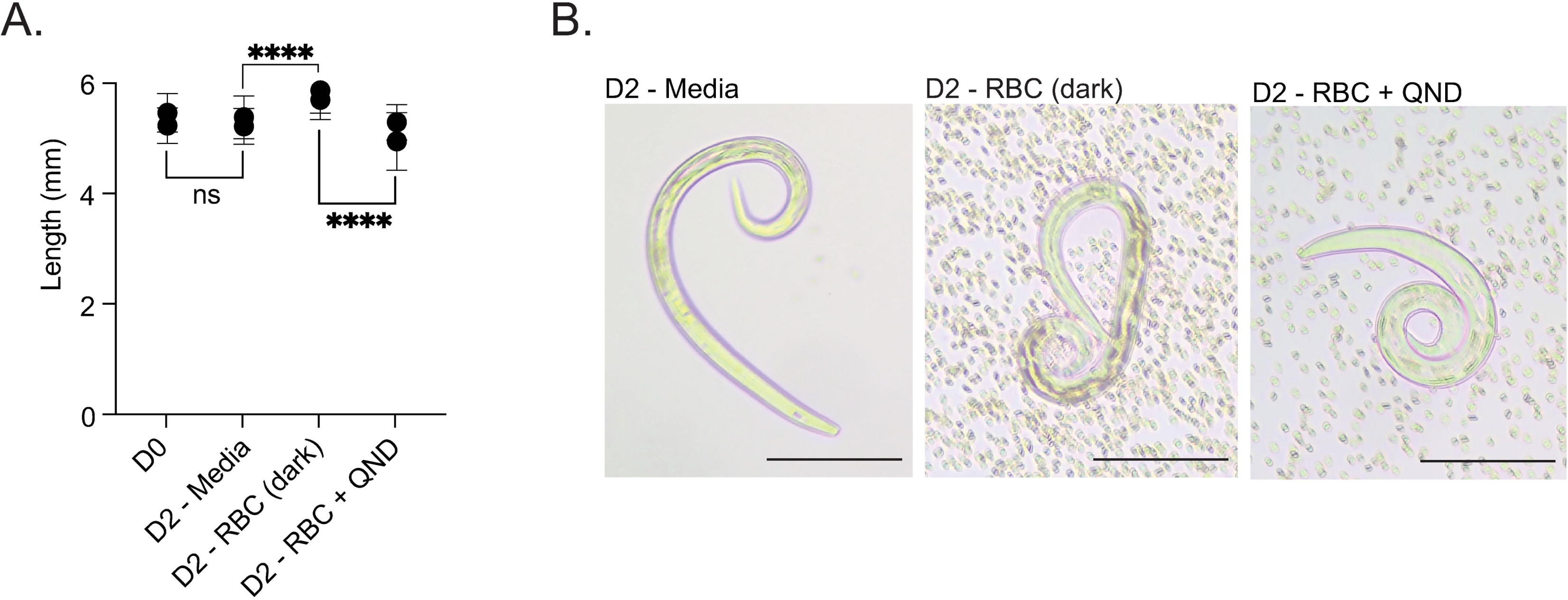
*H. bakeri* L3 growth *in vitro.* Exsheathed larvae were cultured *in vitro* in triplicate with media or 2 x 10^8^ cells per ml murine red blood cells (RBC) for 2 days in the presence or absence of 100 mM QND. A subset of larvae were photographed and measured using ImageJ. (A) The length of larvae. Each dot represents one experiment which is the mean value of 22-64 individual worms dependent on treatment. (B) Photographs of representative exsheathed larvae after culture. N = 3 triplicate wells per group per experiment, 2 independent experiments, Two-way ANOVA, **** = p<0.0001.

At D5 and D7, *H. bakeri* tissue developing stages are relatively easily identifiable by eye as worms with light or dark intestine (LI vs. DI) depending on their pigmentation (Sup Fig 3A). We isolated D5 worms, sexed and sorted them as LI or DI. We cultured the worms *in vitro* for 48 hours, in the presence/absence of RBCs, and assessed their growth to determine whether a darker intestinal pigmentation (likely due to blood feeding) was associated with a growth advantage. Although there was no initial difference in size between the LI/DI D5 worms, both the DI female and male worms grew longer in the presence of RBCs than those in media, while the LI worms did not (Table 2). Both DI and LI worms were ingesting RBCs as demonstrated by the change in their intestinal pigmentation (Sup Fig 3B). When recreating this scenario *in vivo* by measuring freshly isolated worms at D7, not only were there no growth differences between either male or female LI vs. DI worms, but worms were also significantly longer than those of the same age cultured *in vitro* (Table 2). Unlike with the D5 larval worms, when culturing the D7 immature adult worms for three days, we did not observe a growth difference between the LI vs DI worms in the presence/absence of RBCs for either female or male worms (Table 2). The D7 worms grew very little, if at all, *in vitro*: their length after three days of culture was not different to their length at isolation, so growth may not be the most adequate parameter to assess at this time point.

**Table 2:**
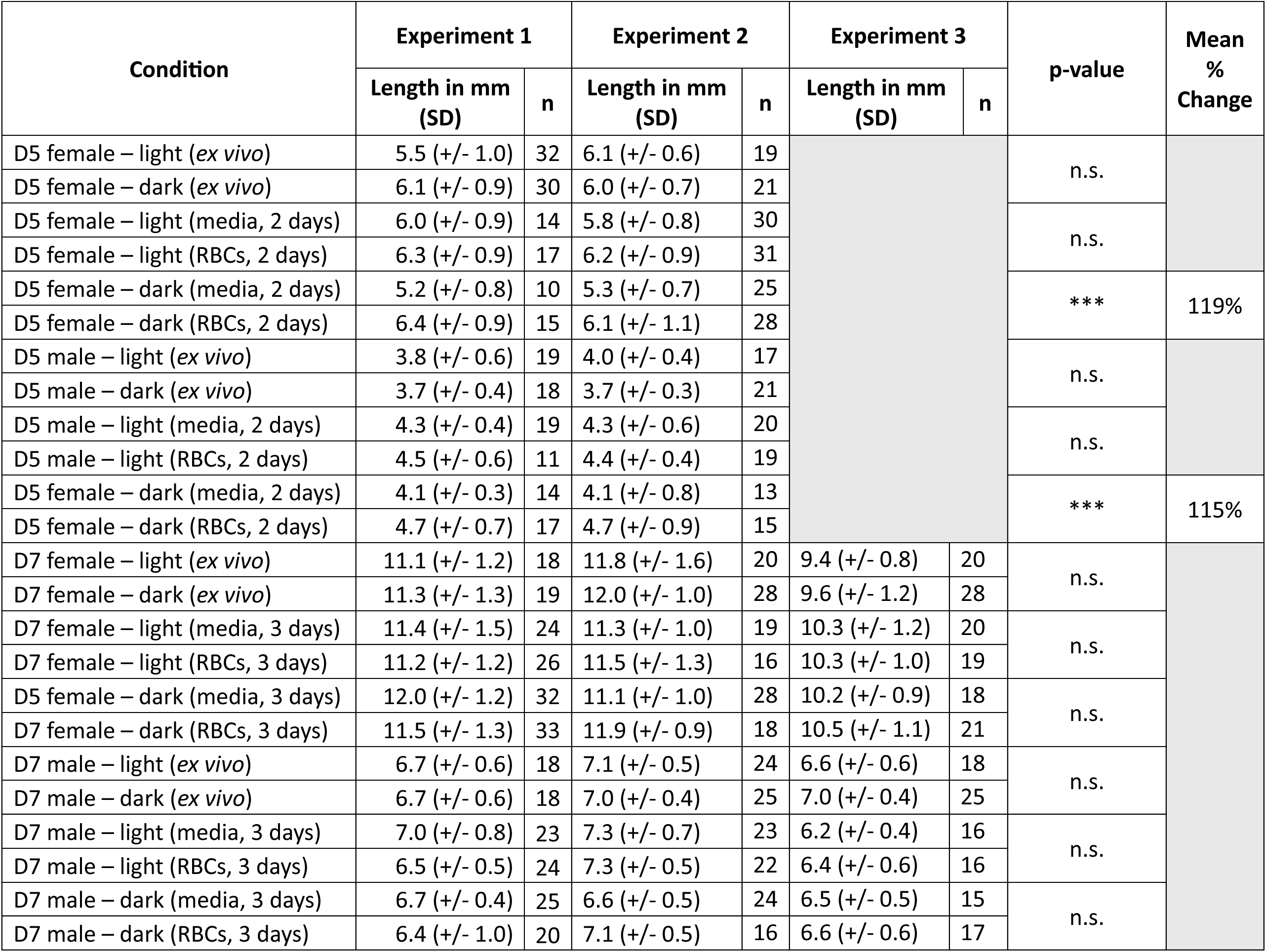
*H. bakeri* blood feeding in developing stages. C57Bl/6 mice were infected orally with 400 infective L3s. At D5 and D7, worms were removed from the intestinal mucosa, sexed and identified as either LI or DI based on pigment. Worms were placed in culture with either media or 2 x 10^8^ cells per ml murine RBC. After two (D5) and three (D7) days of incubation, a subset (minimum of 10 and maximum of 33 individual worms per experiment) of female and male worms were photographed and measured using ImageJ. Two-way ANOVA, n.s. = non significant, *** = p<0.001.

### *In vivo,* DI *H. bakeri* tissue dwelling stages have increased gene expression of blood feeding associated genes

To assess whether we could detect blood feeding in developing worms via a transcriptomic signature, we isolated and sexed D6 and D7 worms with dark and light intestines (DI and LI) for bulk RNA transcriptomics. The worms clustered separately based on transcriptome wide differential expression (Fig 5A). That worms more closely cluster to their developmental time point (D6 DI with D6 LI and D6 DI with D7 LI) confirms that the worms are synchronised and that the darkness of the worms’ intestine was not simply due to one group being ‘older’ (Sup Figure 4).

**Figure 5:**
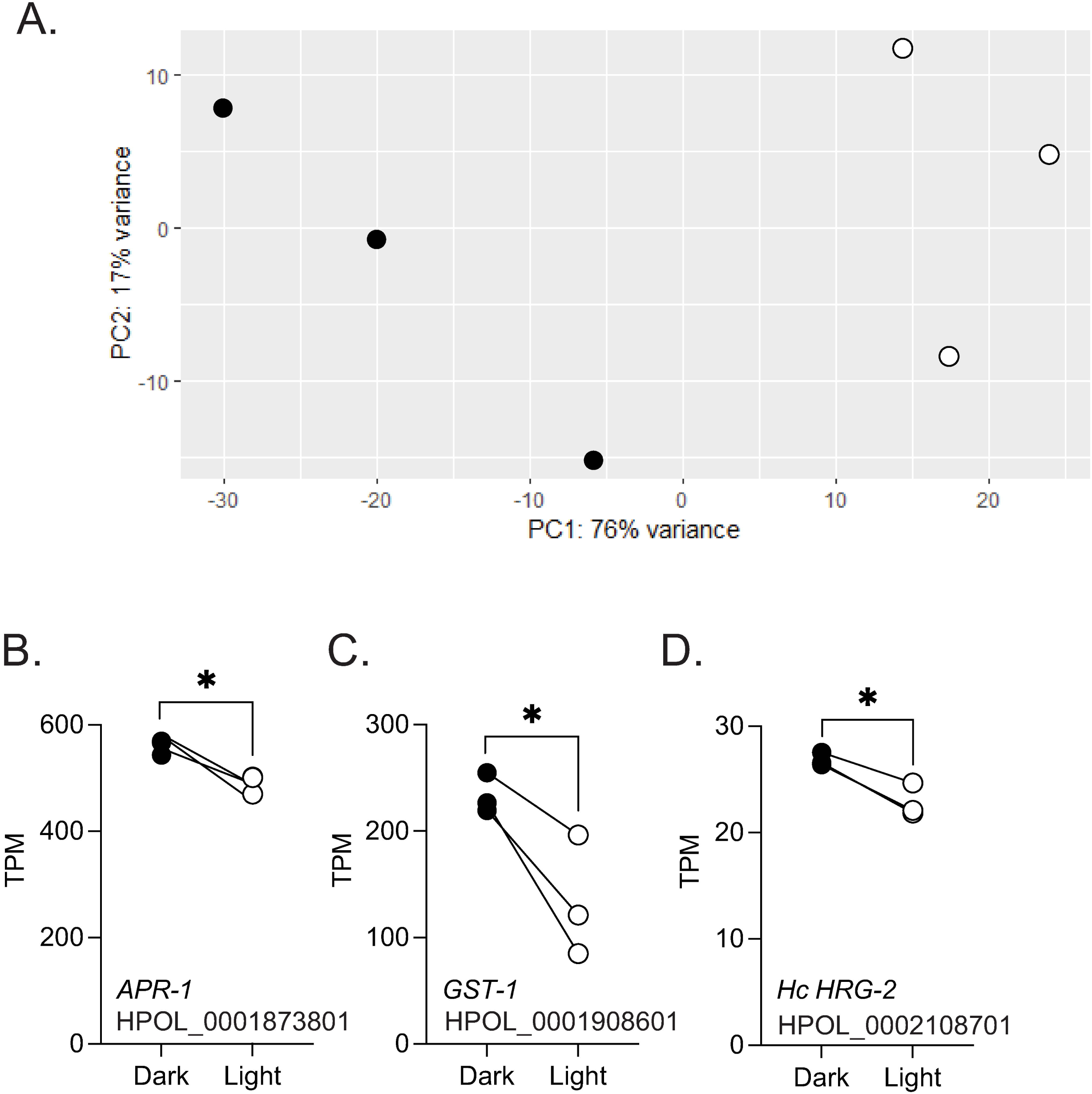
Gene expression of DI vs. LI D7 *H. bakeri* parasitic stages. C57Bl/6 mice were infected orally with 400 infective L3s. At D7, worms were removed from the intestinal tissue and identified as LI or DI for bulk RNA transcriptomics analysis. The experiment was repeated three times. (A) PCA biplot of transcriptomes from DI vs LI worms. Gene expression levels (TPM) of the *APR-1* (B), *GST-1* (C) and *HRG-2* (D) orthologues in *H. bakeri*. N = 3 independent worm collections per time point.

We identified *H. bakeri* orthologues to genes associated with nematode blood-feeding: HPOL_0001873801 for *Na*-APR1; HPOL_0001827001 and HPOL_0001827201 as co-orthologues of *Na*-APR2; a lineage specific expansion of HPOL_0001908601/501/401 and HPOL_0002218401 for *Ac*-GST-1; and HPOL_0001315501 for *Ac*-MEP-1. We could not identify an orthologue for *Ac*-CP-2 in *H. bakeri*.

When we measured their expression within the DI vs. LI *H. bakeri* worms, we found the expression of the *Hb*-APR-1 and one of the *Hb*-GST-1s to be increased in DI worms; this was not the case for *Hb-*APR-2s nor in *Hb*-MEP-1. (Fig 5B and 5C, Sup Fig 5A and 5B).

With regards to the HRGs, we find that neither of the *H. bakeri* orthologues to *Bm*-HRG1 (HPOL_00001917901) or *Bm*-HRG2 (HPOL_000497501) are decreased in the DI worms and that only one of the two orthologues to *Hc*-HRG2 (HPOL_0002148701) is increased in DI worms (Fig 5D, Sup Fig 5D).

### *In vivo,* DI *H. bakeri* tissue dwelling stages differ in their globin and collagen gene expression

When focusing on the potential impact of *in vivo* blood feeding on D7 worms, we first looked at the most expressed genes in both DI and LI worms. In the top 25 genes expressed by either LI or DI worms, 20 were expressed by both (Table 3). The genes in this list comprise 9 globins and 8 collagens. While the exact function of these genes is unknown, globins are involved in O2 transport [25] and collagens in cuticle building [26], both of which have been implicated in worm development, moulting and growth. Of these top 20 genes, five were significantly increased in the DI worms including two glb-1 orthologues (HPOL_0000347301 and HPOL_0001997101) and a col-122 orthologue (cuticle collagen in *C. elegans*, HPOL_0002127501).

**Table 3:**
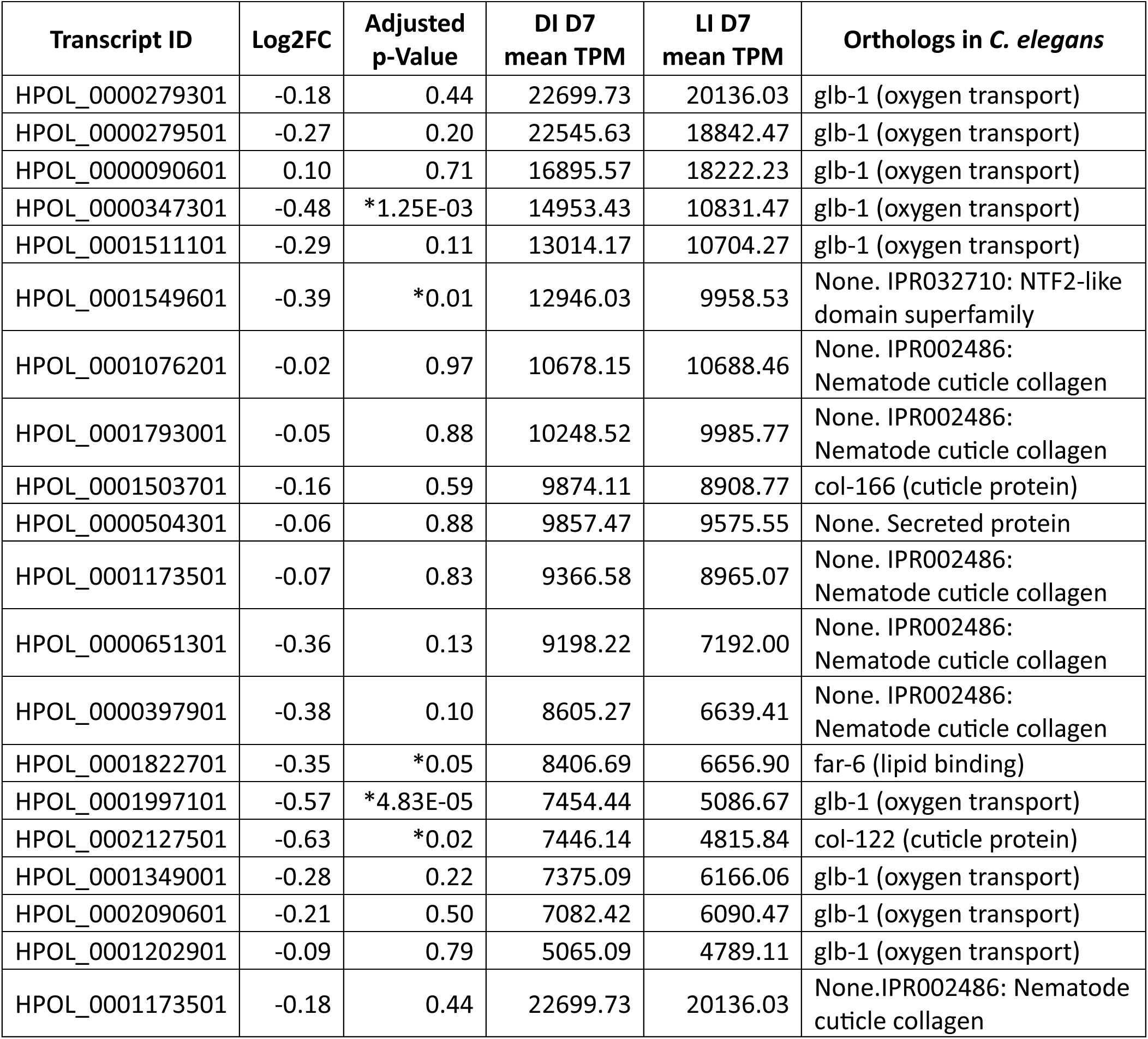
Transcriptomic differences in highly differentially expressed collagen genes in DI vs LI tissue dwelling *H. bakeri.* C57Bl/6 mice were infected orally with 400 infective L3s. At D7, worms were removed from the intestinal tissue and identified as DI or LI for bulk RNA transcriptomics analysis. The experiment was repeated three times. Table shows differences with a Log2FC>-4.

When focusing on the genes that had a fold change of at least 16 (padj<0.05; log2FC≥4) between the DI and LI worms, we obtained 22 genes. Of these, ten genes were cuticle collagen genes (IPR002486: Nematode cuticle collagen, Table 4) and 11 genes had no known function (Sup Table 1). Three genes had orthologues in *C. elegans* and the expression patterns of all *C. elegans* orthologues was consistent with larval expression. When comparing the expression profile of the collagen genes at different lifecycle stages using our previously published transcriptomic data [20], we confirmed that these genes were expressed at the tissue dwelling growing stages (D5 and D7), but not during the adult stages (D10 and D21, Sup Fig 4B).

**Table 4:** Transcriptomic expression of *C. elegans glb-1* orthologues in DI and LI tissue dwelling *H. bakeri.* C57Bl/6 mice were infected orally with 400 infective L3s. At D7, worms were removed from the intestinal tissue and identified as DI or LI for bulk RNA transcriptomics analysis. The experiment was repeated three times. Table shows differences with an adjusted p-value< 0.05.

| Transcript ID | Log2FC | Adjusted p-Value | DI (TPM) | LI (TPM) | Annotation | Orthologs (species #) | Origin |
| --- | --- | --- | --- | --- | --- | --- | --- |
| HPOL_0001529701-mRNA-1 | -10.73 | 2.90E-45 | 324.03 | 0.19 | IPR002486:Nematode cuticle collagen | 48 | Clade 1, 3, 4, 5 |
| HPOL_0002140101-mRNA-1 | -9.83 | 1.78E-38 | 98.79 | 0.11 | IPR002486:Nematode cuticle collagen | 49 | Clade 1, 3, 4, 5 |
| HPOL_0001168501-mRNA-1 | -6.44 | 8.11E-31 | 213.08 | 2.43 | IPR002486:Nematode cuticle collagen | 11 | Clade 3, 4, 5 |
| HPOL_0000732501-mRNA-1 | -5.99 | 3.44E-13 | 303.40 | 4.65 | IPR002486:Nematode cuticle collagen | 48 | Clade 1, 3, 4, 5 |
| HPOL_0002134001-mRNA-1 | -5.48 | 7.44E-41 | 4.24 | 0.10 | IPR002486:Nematode cuticle collagen. Orthologue to *C. elegans col-79* where highly expressed in L4 stage. | 61 | Clade 3, 4, 5 |
| HPOL_0000722101-mRNA-1 | -5.44 | 6.33E-18 | 281.85 | 6.38 | Orthologue to *C. elegans col-151*, where, highly expressed in L4 stage. KO is lethal. | 93 | Clade 1, 3, 4, 5 |
| HPOL_0001071801-mRNA-1 | -5.31 | 8.90E-23 | 68.81 | 1.71 | IPR002486:Nematode cuticle collagen. Orthologue to *C. elegans col-107*, where most highly expressed in L4 stage. | 83 | Clade 3, 4, 5 |
| HPOL_0001089301-mRNA-1 | -4.83 | 2.00E-03 | 1674.98 | 56.05 | IPR002486:Nematode cuticle collagen | 52 | Clade 3, 4, 5 |
| HPOL_0000384201-mRNA-1 | -4.43 | 3.88E-04 | 6.40 | 0.31 | IPR002486:Nematode cuticle collagen | 37 | Clade 3, 4, 5 |
| HPOL_0000176601-mRNA-1 | -4.11 | 4.01E-10 | 8.67 | 0.48 | IPR002486:Nematode cuticle collagen | 33 | Clade 3, 4, 5 |

When looking at the *C. elegans glb-1* (WBGene00014030) orthologues, of which there are 17 in *H. bakeri* (Table 5), 4 were differentially expressed in the DI vs LI D7 worms. Of those four, the three with the highest expression were increased in DI vs. LI and the most lowly expressed has a higher expression in LI vs. DI worms.

**Table 5.**

| Transcript ID | Log2FC | Adjusted p-Value | DI (TPM) | LI (TPM) |
| --- | --- | --- | --- | --- |
| HPOL_0000279301-mRNA-1 | -0.18 | 0.44 | 22699.73 | 20136.03 |
| HPOL_0000279501-mRNA-1 | -0.27 | 0.20 | 22545.63 | 18842.47 |
| HPOL_0000090601-mRNA-1 | 0.10 | 0.71 | 16895.57 | 18222.23 |
| HPOL_0000347301-mRNA-1 | -0.48 | *1.6E-3 | 14953.43 | 10831.47 |
| HPOL_0001511101-mRNA-1 | -0.29 | 0.11 | 13014.17 | 10704.27 |
| HPOL_0001997101-mRNA-1 | -0.57 | *3.4E-3 | 7454.44 | 5086.67 |
| HPOL_0001349001-mRNA-1 | -0.28 | 0.22 | 7375.09 | 6166.06 |
| HPOL_0002090601-mRNA-1 | -0.21 | 0.50 | 7082.42 | 6090.47 |
| HPOL_0001202901-mRNA-1 | -0.09 | 0.79 | 5065.09 | 4789.11 |
| HPOL_0001595301-mRNA-1 | -0.41 | *0.01 | 4947.92 | 3745.37 |
| HPOL_0001203201-mRNA-1 | 0.00 | 1.00 | 4010.60 | 4030.78 |
| HPOL_0001810101-mRNA-1 | -0.09 | 0.76 | 3146.61 | 2974.51 |
| HPOL_0001203301-mRNA-1 | 0.21 | 0.55 | 2939.60 | 3428.67 |
| HPOL_0002061901-mRNA-1 | -0.20 | 0.35 | 2453.66 | 2172.59 |
| HPOL_0001203101-mRNA-1 | 0.67 | *0.03 | 1606.76 | 2578.81 |
| HPOL_0001434001-mRNA-1 | -0.18 | 0.66 | 603.45 | 531.34 |
| HPOL_0000090501-mRNA-1 | -0.27 | 0.07 | 89.18 | 75.00 |

## DISCUSSION

Haem, a complex of iron with protoporphyrin IX, is essential for the function of all aerobic cells. It plays key roles in oxidative metabolism, cellular differentiation, gene transcription and protein translation/stability [27]. However, nematodes have lost the ability to synthesise haem *de novo* and cannot catabolise haem using haem oxygenase [2]. Many parasitic nematodes obtain haem from their hosts, as dietary haem is more easily absorbed than inorganic iron.

Consequently, obligate blood-feeding has evolved independently multiple times independently in nematodes, particularly in the Strongylida. Notable examples include the barber pole worm, *H. contortus*, and hookworms, *N. americanus* and *A. caninum*, all of which have specialised feeding structures and secrete anti-coagulants to access and maintain their food source [28–32]. It remains uncertain as to whether the hookworms (Ancylostomatoidea) are monophyletic with phylogenetics based on mitochondrial genomes and 18S RNA genes pointing to paraphyly [33–36]. Regardless of the phylogenetic relationships, all these worms feed exclusively on blood and induce chronic anemia in their hosts [37,38].

*Heligmosomoides bakeri*, another nematode in the Strongylida, is typically classified as a non-blood feeder. However, its larvae and immature adults are encysted within the host intestinal tissue, causing damage and positioning themselves close to host blood. Here, we have shown that both larvae and adults can ingest blood *in vitro*, which provides developing worms with a growth advantage. Meanwhile *in vivo*, we observed worms with dark and light intestines (DI and LI), indicative of differences in blood ingestion. Despite no significant size differences between DI and LI worms, several cuticle collagens—key components of growth—were more highly expressed in DI worms. We also observed differential expression of *H. bakeri* globin genes. *H. bakeri* has 17 co-orthologues of *Ce-glb-1*; three of the top ten were upregulated in DI worms. While the exact function of *Ce-glb-1* is not fully understood, it has been associated with O_2_ binding, NO scavenging or O_2_ storage [39]. Although DI worms exhibited distinct expression of a subset of *glb-1* isoforms, the expression of other *H. bakeri glb-1* co-orthologues remained high in both DI and LI worms, with four of these genes being extremely highly expressed (TPMs>10000). This suggests that the darker red coloration seen in DI worms is not solely attributable to the presence of globins, as might be expected if these worms were more developed. Furthermore, when clustering the worms by the number of days post-infection (D6 vs D7), rather than by intestinal coloration (dark vs light), we observed that the worms were synchronized by age.

An advantage of blood feeding may be faster development. For example, DI D7 worms may have moulted or be ready to moult sooner than the LI worms, as collagen expression and availability of oxygen are critical for this process [20,26]. Our understanding of developmental cues, such as moulting, in parasitic nematodes remains limited. Many genes have been implicated in *C. elegans* moulting, however, some like *Ce-col-12*, *Ce-sqt-1*, and *Ce-dpy-13*, are absent in *H. bakeri* [40,41]. While it is possible, their absence is due to errors in the *H. bakeri* genome assembly or annotation [42], we note that they are also absent from the assemblies of *N. brasiliensis* and *H. contortus*, suggesting a divergence in the inventory of genes involved in moulting. The pig nodule worm, *Oesophagostomum dentatum*, has been proposed as a model for studying moulting in parasitic nematodes [43]. Although many of the moulting-associated genes in *O. dentatum* do have orthologues in *H. bakeri*, the temporal expression differences between L4 and adult have not been full characterized, making it difficult to assess their relevance in other species.

Somewhat surprisingly, blood feeding *in vitro*, though leading to darker pigmentation and growth advantages, did not result in measurable size differences *in vivo*. This could indicate that tissue-dwelling worms may not frequently encounter RBCs or that the amount of blood ingestion varies among individual worms. Furthermore, the substantial growth observed between D5 and D7 worms *in vivo*, with minimal growth *in vitro*, suggests that current *in vitro* conditions do not accurately reflect the *in vivo* environment. The D5 parasites could be using oxygen from the RBC incubation to prepare for their next moult, but without understanding the exact moulting triggers, this effect cannot be adequately measured. Similarly, D7 worms showed no growth *in vitro*, which may indicate they are preparing for tissue exit, a process for which we also lack environmental cues.

*Nippostrongylus brasiliensis*, a close relative *H. bakeri*, has a lifecycle similar to hookworms; it enters its murine host through the skin, migrates to the lung, and then moves to the gut, where it undergoes a final moult. Unlike the hookworms, it does not have specialised feeding structures [15]. To our knowledge, no anti-coagulants have been identified for *N. brasiliensis*, and we could not find homologs to known anti-coagulants from *Ancylostoma*, *Necator*, or *Haemonchus* (Rich and Wasmuth *pers. comms.*). Although a new anti-coagulant was recently identified for *Ascaris suum*—a more distantly related nematode but one whose larvae also traverse through the lungs—its function was linked to the migration and immunomodulation rather than blood feeding [44].

Further, support for *N. brasiliensis* as an obligate blood feeder relies, in large part on the observation that the worm causes anemia and the claim that the use of quinolone drugs blocks a haem detoxification pathway [13]. Administration of these drugs reduced parasite fitness and improved anemia in infected mice. We also observed that larval stages of *H. bakeri* caused a small but significant anemia. Furthermore, administration of quinidine blocked uptake of RBC by *H. bakeri*. However, a bead-feeding assay demonstrates that quinidine blocks feeding through the suppression of pharyngeal pumping, consistent with findings in *C. elegans* [24].

These and other observations lead us to make two conclusions. First, both *N. brasiliensis* and *H. bakeri* larvae and immature adults feed on host blood to acquire nutrients and the oxygen necessary for growth and moulting. Second, the adults of both species—residing in the host’s intestinal lumen—do not have easy access to blood and instead graze on host tissue, which is consistent with previous findings [14,18].

The requirement for host blood among parasitic nematodes exists on a spectrum. Some species, for example the microfilariae of *B. malayi*, spend significant time in circulation. Other species, such as the larvae *Trichinella*, *Ascaris*, *Strongyloides* and *Nippostrongylus*, migrate to different host organs through the circulatory system. The larvae of *Anisakis*, *Trichuris* and *Heligmosomoides* damage tissue which may result in broken blood vessels and the leak of red blood cells which could be a source of food for tissue dwelling stages. Finally, the adults of *Haemonchus*, *Necator*, *Ancylostoma*—and other hookworms—actively seek blood, puncturing or ripping the host’s intestinal lining. Even the free-living *C. elegans* can survive on a diet of RBC [12]; this ability to acquire nutrition from variable sources provides another putative preadaptation towards parasitism. We would describe *H. bakeri* as a metabolic opportunist [45], feeding on blood if and when available.

## ACKNOWLEDGEMENTS

We would like to thank the University of Calgary Animal Facility technicians, especially Dawn Martin for her help, as well as Dr Carrie Shemanko and Wic Wildering for the use of their microscopes. We also acknowledge the investment in high-performance computing resources by the Faculty of Veterinary Medicine and Research Computing at the University of Calgary.

## DATA ACCESS

All data not available in the manuscript or supplemental materials are available from the corresponding author upon request.

---

**Supplementary Figure 1: Quinidine (QND) inhibits feeding in *H. bakeri* larvae.** Exsheathed *H. bakeri* larvae were incubated with fluorescent beads of various sizes (0.5 μm and 6μm) *in vitro*, after 5 days ingestion of beads was assessed in the presence or absence of QND. (A) % of exsheathed larvae having ingested beads. Experiments were repeated twice with a minimum of 39 and a maximum of 142 larvae per well. (B) Representative images of fluorescent beads detected inside or outside of larvae. (C) Fluorescence intensity of larvae ingested beads incubated in media or media with QND. Experiment was performed once, with a minimum of 5 and a maximum of 14 larvae per well. (D) Representative images of fluorescence inside worms with or without beads or QND.

**Supplementary Figure 2: Quinidine (QND) inhibits feeding in *H. bakeri* adult worms.** C57Bl/6 mice were infected orally with 200 infective L3s. At D10, worms were removed from the intestinal tissue and were incubated with fluorescently labelled beads of various sizes (0.5 μm and 6 μm) *in vitro*, after 5 days ingestion of beads was assessed in the presence or absence of QND. (A) % of adult worms ingested beads. Experiment was performed twice, with a minimum of 8 and a maximum of 20 adults per well. (B) Representative images of fluorescent beads detected inside adult worms. (C) Fluorescence intensity of adult worms ingested 0.5 μm beads incubated in media or media with QND. Experiment was performed once, with a minimum of 3 and a maximum of 12 adults per well. (D) Representative images of fluorescence inside worms with or without beads or QND after ingestion of 0.5 μm beads. (E) Fluorescence intensity of adult worms ingested 6 μm beads incubated in media or media with QND. Experiment was performed once, with a minimum of 7 and a maximum of 18 adults per well. (F) Representative images of fluorescence inside worms with or without beads or QND after ingestion of 6 μm beads.

**Supplementary Figure 3: Tissue stage *H. bakeri* pigmentation.** (A) Representative photographs of DI vs. LI D7 worms taken *in vivo* from the outside of the intestine. (B) Representative photographs of DI vs. LI D5 worms taken from isolated worms which were placed on a slide and covered with a coverslip immediately after being isolated or after 2 days of culture with RBCs.

**Supplementary Figure 4: Transcriptomic profiles of developing *H. bakeri.*** Bulk RNA from *H. bakeri* worms was extracted at two different time points: D6 and D7 for RNA sequencing. C57Bl/6 mice were infected orally with 400 infective L3s. At D6, worms were removed from the intestinal tissue and identified as LI or DI for bulk RNA transcriptomics analysis. PCA biplot of transcriptomes from DI vs LI worms at D6 (single sample) and D7 (in triplicate).

**Supplementary Figure 5: Gene expression of blood feeding associated genes in *H. bakeri.*** C57Bl/6 mice were infected orally with 400 infective L3s. At D7, worms were removed from the intestinal tissue and identified as LI or DI for bulk RNA transcriptomics analysis. Gene expression levels (TPM) of the *APR-2* (A), *GST-1* (B), *MEP-1* (C) and *HRG* (D) orthologues in *H. bakeri*. N = 3 independent worm collections per time point.

**Supplementary Table 1: Transcriptomic differences in highly differentially expressed genes with no known function in DI vs LI tissue dwelling *H. bakeri.*** C57Bl/6 mice were infected orally with 400 infective L3s. At D7, worms were removed from the intestinal tissue and identified as LI or DI for bulk RNA transcriptomics analysis. The experiment was repeated three times. Table shows differences with a Log2FC>-4.

## REFERENCES

1. Toh SQ, Gobert GN, Malagón Martínez D, Jones MK. Haem uptake is essential for egg production in the haematophagous blood fluke of humans, *Schistosoma mansoni*. The FEBS Journal. 2015;282: 3632–3646. doi:10.1111/febs.13368

2. Perner J, Gasser RB, Oliveira PL, Kopáček P. Haem Biology in Metazoan Parasites – ‘The Bright Side of Haem.’ Trends in Parasitology. 2019;35: 213–225. doi:10.1016/j.pt.2019.01.001

3. Rajagopal A, Rao AU, Amigo J, Tian M, Upadhyay SK, Hall C, et al. Haem homeostasis is regulated by the conserved and concerted functions of HRG-1 proteins. Nature. 2008;453: 1127–1131. doi:10.1038/nature06934

4. Luck AN, Yuan X, Voronin D, Slatko BE, Hamza I, Foster JM. Heme acquisition in the parasitic filarial nematode *Brugia malayi*. FASEB j. 2016;30: 3501–3514. doi:10.1096/fj.201600603R

5. Zhou J-R, Bu D-R, Zhao X-F, Wu F, Chen X-Q, Shi H-Z, et al. Hc-hrg-2, a glutathione transferase gene, regulates heme homeostasis in the blood-feeding parasitic nematode Haemonchus contortus. Parasites Vectors. 2020;13: 40. doi:10.1186/s13071-020-3911-z

6. Ranjit N, Zhan B, Hamilton B, Stenzel D, Lowther J, Pearson M, et al. Proteolytic degradation of hemoglobin in the intestine of the human hookworm Necator americanus. J Infect Dis. 2009;199: 904–912. doi:10.1086/597048

7. Zhan B, Liu S, Perally S, Xue J, Fujiwara R, Brophy P, et al. Biochemical Characterization and Vaccine Potential of a Heme-Binding Glutathione Transferase from the Adult Hookworm Ancylostoma caninum. Infection and Immunity. 2005;73: 6903–6911. doi:10.1128/iai.73.10.6903-6911.2005

8. van Rossum AJ, Jefferies JR, Rijsewijk FAM, LaCourse EJ, Teesdale-Spittle P, Barrett J, et al. Binding of Hematin by a New Class of Glutathione Transferase from the Blood-Feeding Parasitic Nematode Haemonchus contortus. Infect Immun. 2004;72: 2780–2790. doi:10.1128/IAI.72.5.2780-2790.2004

9. Williamson AL, Brindley PJ, Abbenante G, Prociv P, Berry C, Girdwood K, et al. Cleavage of hemoglobin by hookworm cathepsin D aspartic proteases and its potential contribution to host specificity. The FASEB Journal. 2002;16: 1458–1460.

10. Diemert DJ, Lobato L, Styczynski A, Zumer M, Soares A, Gazzinelli MF. A Comparison of the Quality of Informed Consent for Clinical Trials of an Experimental Hookworm Vaccine Conducted in Developed and Developing Countries. PLoS Negl Trop Dis. 2017;11: e0005327. doi:10.1371/journal.pntd.0005327

11. Diemert DJ, Zumer M, Campbell D, Grahek S, Li G, Peng J, et al. Safety and immunogenicity of the Na-APR-1 hookworm vaccine in infection-naïve adults. Vaccine. 2022;40: 6084– 6092. doi:10.1016/j.vaccine.2022.09.017

12. Chauhan VM, Pritchard DI. Haematophagic Caenorhabditis elegans. Parasitology. 2019;146: 314–320. doi:10.1017/S0031182018001518

13. Bouchery T, Filbey K, Shepherd A, Chandler J, Patel D, Schmidt A, et al. A novel blood-feeding detoxification pathway in Nippostrongylus brasiliensis L3 reveals a potential checkpoint for arresting hookworm development. Selkirk ME, editor. PLoS Pathog. 2018;14: e1006931. doi:10.1371/journal.ppat.1006931

14. Bansemir AD, Sukhdeo MVK. Intestinal Distribution of Worms and Host Ingesta in Nippostrongylus brasiliensis. The Journal of Parasitology. 2001;87: 1470–1472. doi:10.2307/3285321

15. Seesee FM, Wescott RB, Gorham JR. Visualization of Nippostrongylus brasiliensis by Scanning Electron Microscopy. The Journal of Parasitology. 1977;63: 1135–1137. doi:10.2307/3279871

16. Stevens L, Martínez-Ugalde I, King E, Wagah M, Absolon D, Bancroft R, et al. Ancient diversity in host-parasite interaction genes in a model parasitic nematode. Nat Commun. 2023;14: 7776. doi:10.1038/s41467-023-43556-w

17. Ehrenford FA. The Life Cycle of Nematospiroides dubius Baylis (Nematoda: Heligmosomidae). The Journal of Parasitology. 1954;40: 480–481. doi:10.2307/3273905

18. Bansemir AD, Sukhdeo MVK. The Food Resource of Adult Heligmosomoides polygyrus in the Small Intestine. The Journal of Parasitology. 1994;80: 24–28. doi:10.2307/3283340

19. Johnston CJC, Robertson E, Harcus Y, Grainger JR, Coakley G, Smyth DJ, et al. Cultivation of Heligmosomoides polygyrus: an immunomodulatory nematode parasite and its secreted products. J Vis Exp. 2015; e52412. doi:10.3791/52412

20. Pollo SMJ, Leon-Coria A, Liu H, Cruces-Gonzalez D, Finney CAM, Wasmuth JD. Transcriptional patterns of sexual dimorphism and in host developmental programs in the model parasitic nematode Heligmosomoides bakeri. Parasites Vectors. 2023;16: 171. doi:10.1186/s13071-023-05785-2

21. Dobin A, Davis CA, Schlesinger F, Drenkow J, Zaleski C, Jha S, et al. STAR: ultrafast universal RNA-seq aligner. Bioinformatics. 2013;29: 15–21. doi:10.1093/bioinformatics/bts635

22. Liao Y, Smyth GK, Shi W. featureCounts: an efficient general purpose program for assigning sequence reads to genomic features. Bioinformatics. 2014;30: 923–930. doi:10.1093/bioinformatics/btt656

23. Love MI, Huber W, Anders S. Moderated estimation of fold change and dispersion for RNA-seq data with DESeq2. Genome Biology. 2014;15: 550. doi:10.1186/s13059-014-0550-8

24. Franks CJ, Pemberton D, Vinogradova I, Cook A, Walker RJ, Holden-Dye L. Ionic Basis of the Resting Membrane Potential and Action Potential in the Pharyngeal Muscle of Caenorhabditis elegans. Journal of Neurophysiology. 2002;87: 954–961. doi:10.1152/jn.00233.2001

25. Blaxter ML. Nemoglobins: divergent nematode globins. Parasitol Today. 1993;9: 353–360. doi:10.1016/0169-4758(93)90082-q

26. Page AP, Stepek G, Winter AD, Pertab D. Enzymology of the nematode cuticle: A potential drug target? Int J Parasitol Drugs Drug Resist. 2014;4: 133–141. doi:10.1016/j.ijpddr.2014.05.003

27. Ponka P. Cell biology of heme. Am J Med Sci. 1999;318: 241–256. doi:10.1097/00000441-199910000-00004

28. Carroll SM, Robertson TA, Papadimitriou JM, Grove DI. Scanning electron microscopy of Ancylostoma ceylanicum and its site of attachment to the small intestinal mucosa of the dog. Z Parasitenkd. 1985;71: 79–85. doi:10.1007/BF00932921

29. Weise RW. A Light and Electron Microscopic Study of the Dorsal Buccal Lancet of Haemonchus contortus. The Journal of Parasitology. 1977;63: 854–857. doi:10.2307/3279892

30. Uppal HS, Bal MS, Singla LD, Kaur P, Sandhu BS. Morphometric and scanning electron microscopy based identification of Ancylostoma caninum parasites in dog. J Parasit Dis. 2017;41: 517–522. doi:10.1007/s12639-016-0841-y

31. Abuzeid AMI, Zhou X, Huang Y, Li G. Twenty-five-year research progress in hookworm excretory/secretory products. Parasit Vectors. 2020;13: 136. doi:10.1186/s13071-020-04010-8

32. Geldhof P, Knox D. The intestinal contortin structure in Haemonchus contortus: an immobilised anticoagulant? Int J Parasitol. 2008;38: 1579–1588. doi:10.1016/j.ijpara.2008.05.002

33. Ahmed M, Roberts NG, Adediran F, Smythe AB, Kocot KM, Holovachov O. Phylogenomic Analysis of the Phylum Nematoda: Conflicts and Congruences With Morphology, 18S rRNA, and Mitogenomes. Front Ecol Evol. 2022;9. doi:10.3389/fevo.2021.769565

34. Jex AR, Hall RS, Littlewood DTJ, Gasser RB. An integrated pipeline for next-generation sequencing and annotation of mitochondrial genomes. Nucleic Acids Research. 2010;38: 522–533. doi:10.1093/nar/gkp883

35. van Megen H, van der Elsen S, Holterman M, Karssen G, Mooyman P, Bongers T, et al. A phylogenetic tree of nematodes based on about 1200 full-length small subunit ribosomal DNA sequences. Nematology. 2009;11: 927–950. doi:10.1163/156854109X456862

36. Sultana T, Kim J, Lee S-H, Han H, Kim S, Min G-S, et al. Comparative analysis of complete mitochondrial genome sequences confirms independent origins of plant-parasitic nematodes. BMC Evolutionary Biology. 2013;13: 12. doi:10.1186/1471-2148-13-12

37. Clements ACA, Addis Alene K. Global distribution of human hookworm species and differences in their morbidity effects: a systematic review. The Lancet Microbe. 2022;3: e72–e79. doi:10.1016/S2666-5247(21)00181-6

38. Besier RB, Kahn LP, Sargison ND, Van Wyk JA. The Pathophysiology, Ecology and Epidemiology of *Haemonchus contortus* Infection in Small Ruminants. In: Gasser RB, Samson-Himmelstjerna GV, editors. Advances in Parasitology. Academic Press; 2016. pp. 95–143. doi:10.1016/bs.apar.2016.02.022

39. Geuens E, Hoogewijs D, Nardini M, Vinck E, Pesce A, Kiger L, et al. Globin-like proteins in Caenorhabditis elegans: in vivo localization, ligand binding and structural properties. BMC Biochem. 2010;11: 17. doi:10.1186/1471-2091-11-17

40. Johnstone IL. Cuticle collagen genes: expression in Caenorhabditis elegans. Trends in Genetics. 2000;16: 21–27. doi:10.1016/S0168-9525(99)01857-0

41. Johnstone IL, Barry JD. Temporal reiteration of a precise gene expression pattern during nematode development. The EMBO Journal. 1996;15: 3633–3639. doi:10.1002/j.1460-2075.1996.tb00732.x

42. Mariene GM, Wasmuth JD. Genome assembly variation and its implications for gene discovery in nematode species. bioRxiv; 2024. p. 2024.02.26.582167. doi:10.1101/2024.02.26.582167

43. Ondrovics M, Gasser RB, Joachim A. Recent Advances in Elucidating Nematode Moulting – Prospects of Using *Oesophagostomum dentatum* as a Model. In: Rollinson D, Stothard JR, editors. Advances in Parasitology. Academic Press; 2016. pp. 233–264. doi:10.1016/bs.apar.2015.09.001

44. Diosdado A, Simón F, Morchón R, González-Miguel J. Host-Parasite Relationships in Porcine Ascariosis: Anticoagulant Potential of the Third Larval Stage of Ascaris suum as a Possible Survival Mechanism. Animals (Basel). 2021;11: 804. doi:10.3390/ani11030804

45. Bryant V. Growth and respiration throughout the life-cycle of Nematospiroides dubius Baylis (1926) (Nematoda: Heligmosomidae): the parasitic stages. Parasitology. 1974;69: 97–106. doi:10.1017/S0031182000046217

