## Supplementary figures and images for "Evidence of opportunistic blood feeding in the parasitic nematode *Heligmosomoides bakeri*"

### Supplemental Figure 1

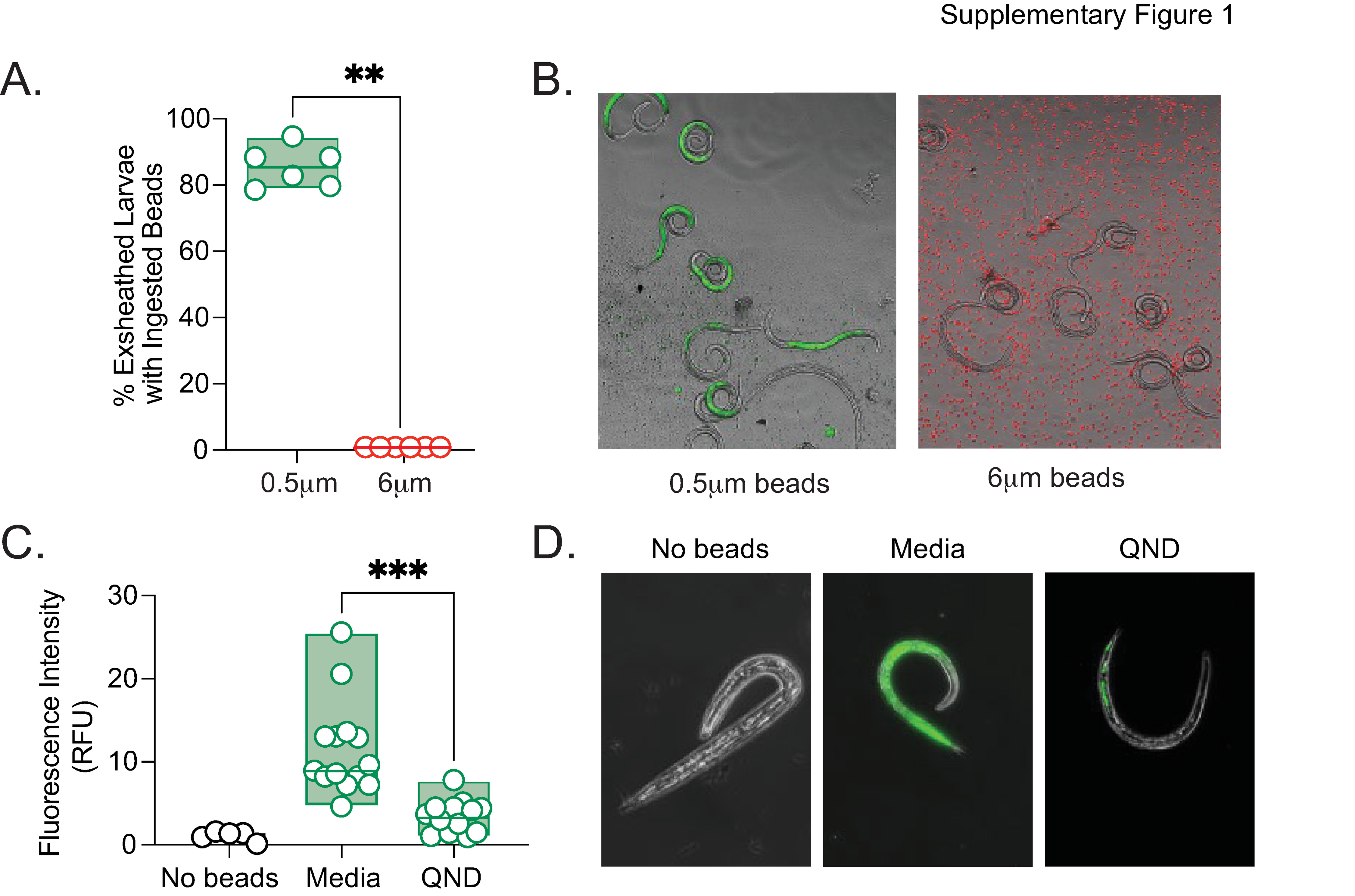

### Supplemental Figure 2

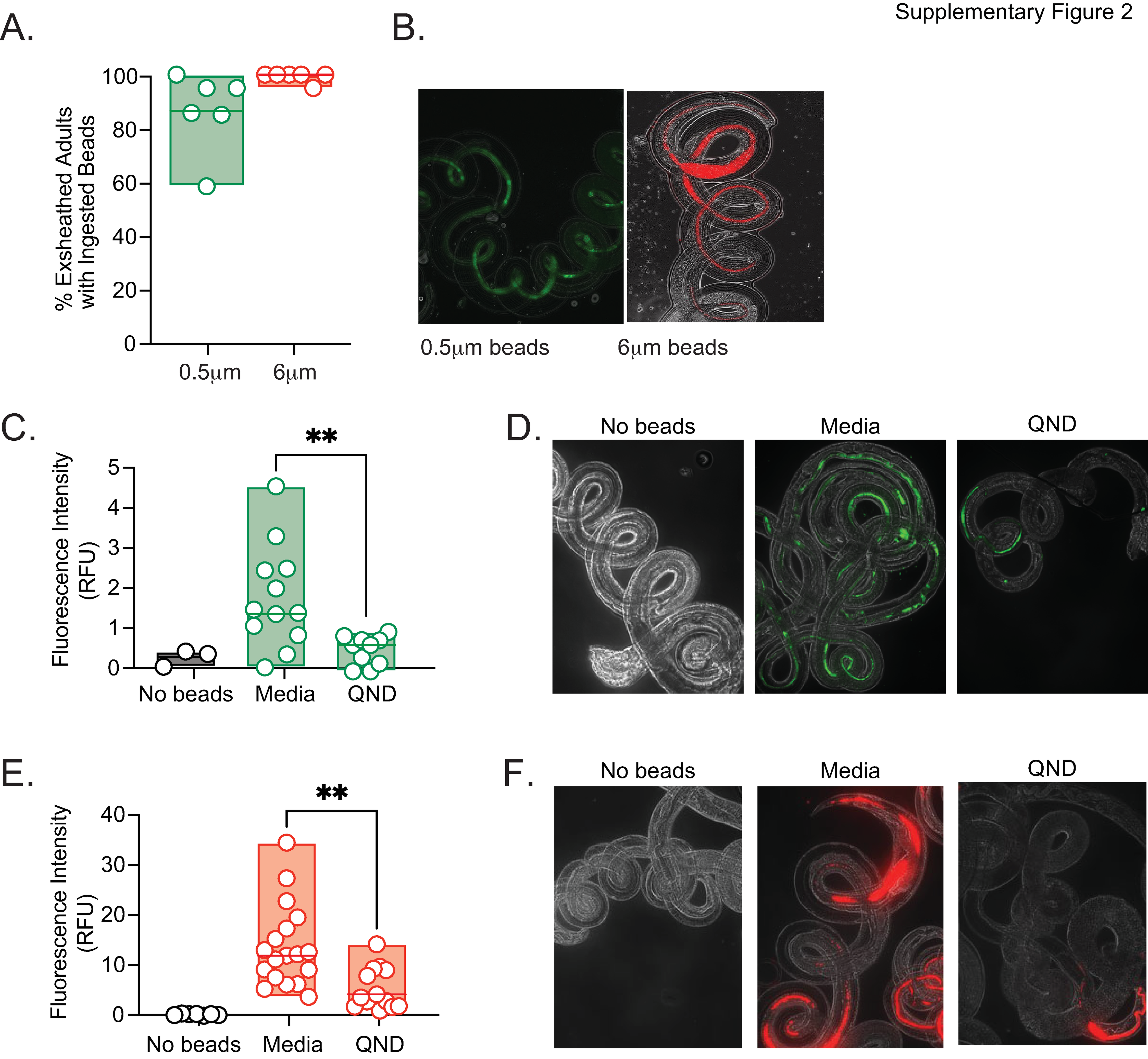

### Supplemental Figure 3

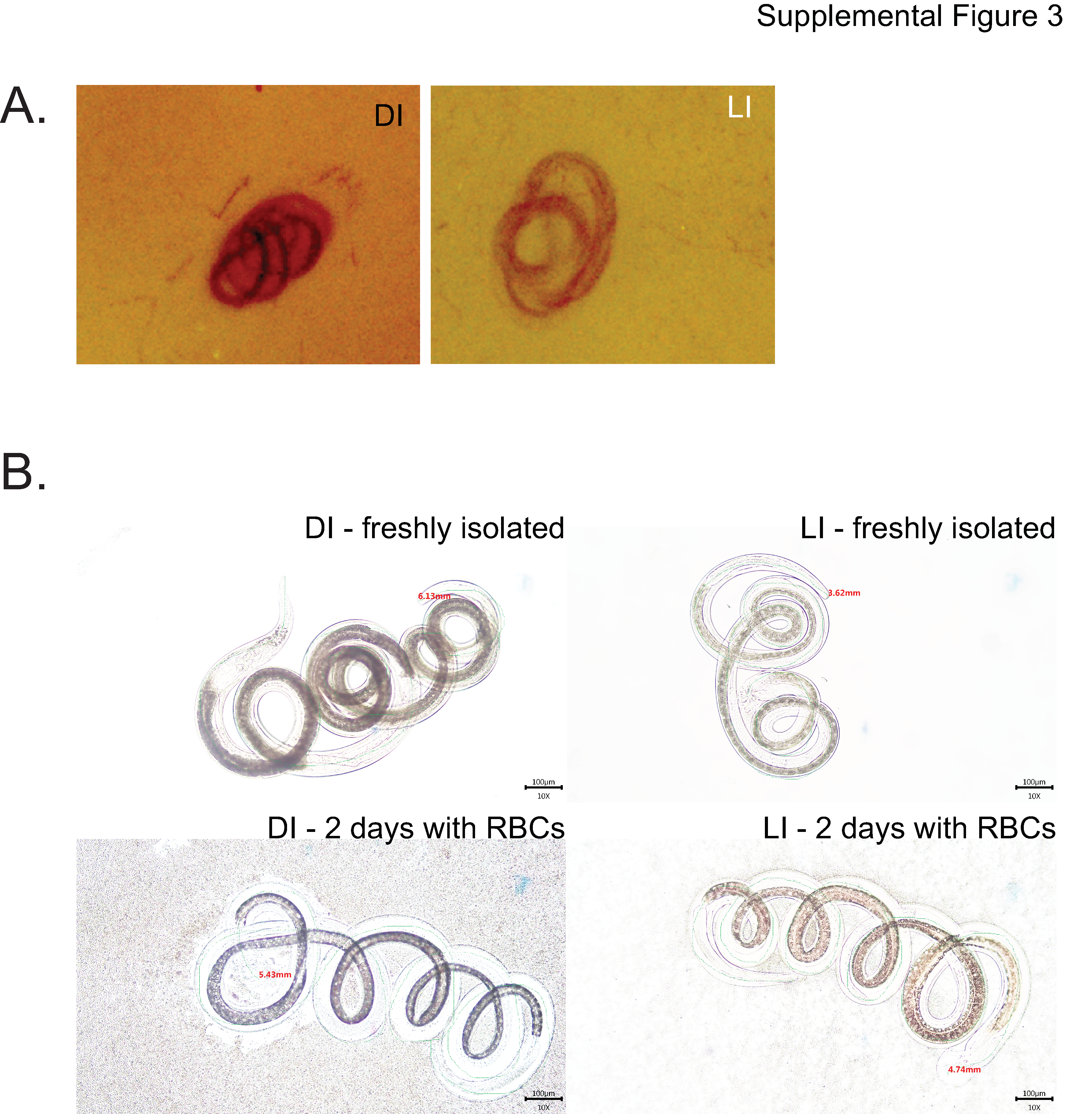

### Supplemental Figure 4

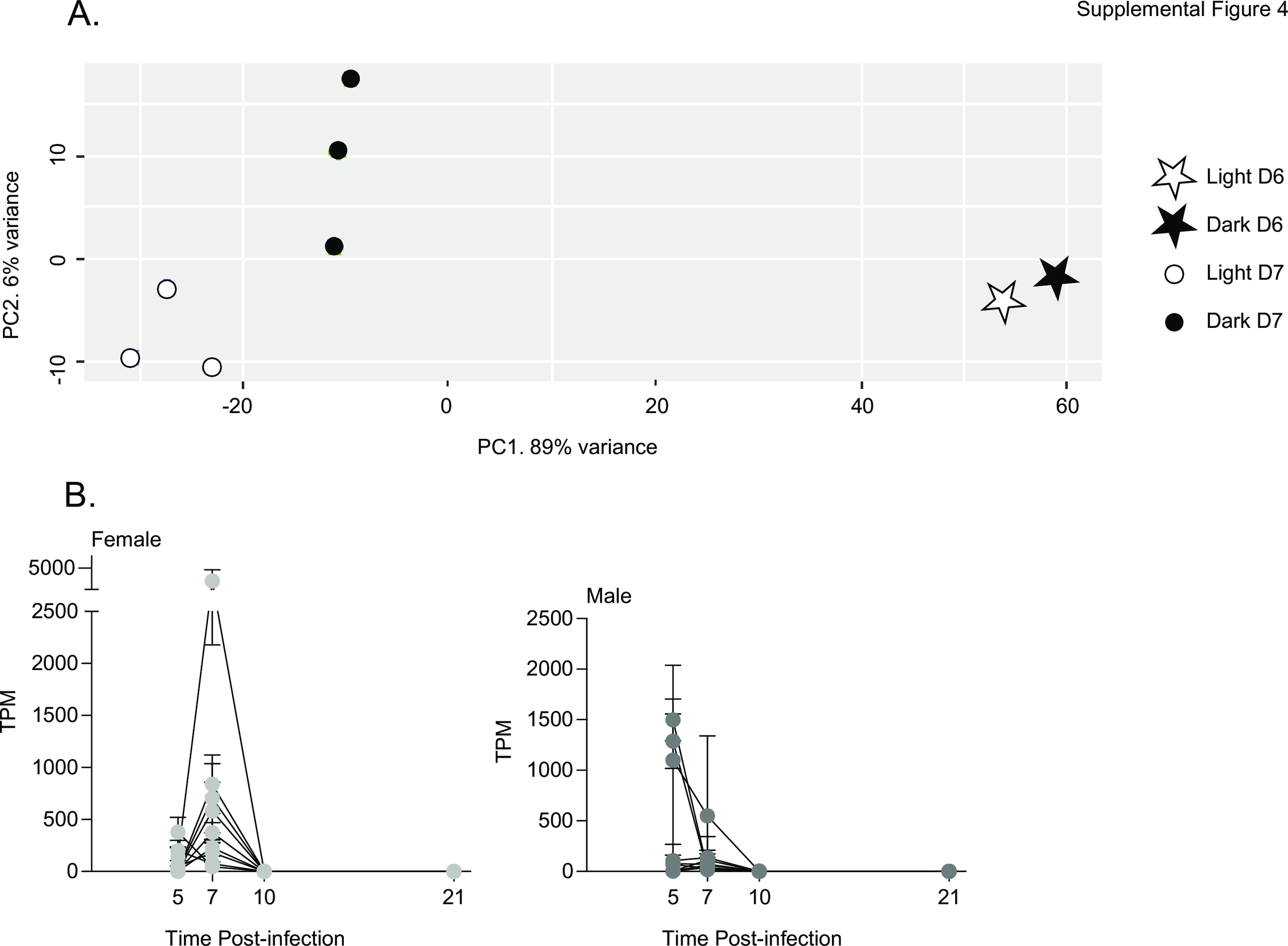

### Supplemental Figure 5

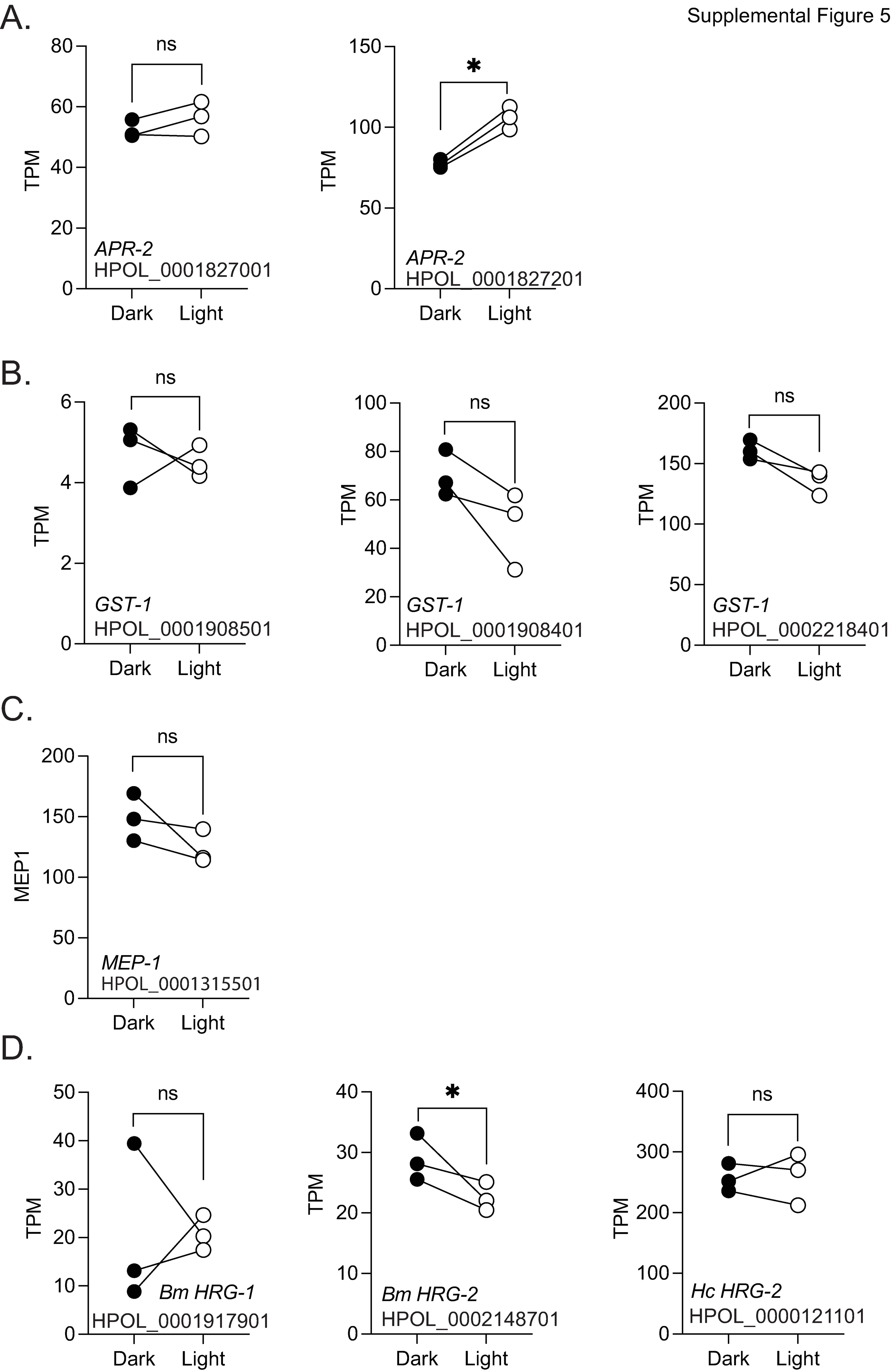
