## Supplemental Table 1 for "Evidence of opportunistic blood feeding in the parasitic nematode *Heligmosomoides bakeri*"

| Transcript ID | Log2FC | Adjusted p-Value | Annotation | Orthologs (species #) | Origin |
|---|---|---|---|---|---|
| HPOL_0002223901-mRNA-1 | -11.22 | 1.50E-13 | NA | None | |
| HPOL_0002363601-mRNA-1 | -9.31 | 6.12E-3 | NA | None | |
| HPOL_0000197701-mRNA-1 | -5.64 | 5.33E-33 | NA | None | |
| HPOL_0001825001-mRNA-1 | -5.31 | 6.14E-36 | IPR035161:Protein of unknown function DUF5332. Orthologue to *C. elegans dct-5,* which is involved in defense response to Gram – bacteria and enriched in larval stages | 46 | Clade 3, 4, 5 |
| HPOL_0000584501-mRNA-1 | -5.31 | 6.54E-19 | Orthologue to *C. elegans Y57G11C.42*. In *C. elegans,* expression higher in larval vs. adult stage. | 112 | Metazoa |
| HPOL_0001398001-mRNA-1 | -4.86 | 1.15E-05 | NA | None | |
| HPOL_0001381801-mRNA-1 | -4.60 | 1.84E-2 | NA | None | |
| HPOL_0000146901-mRNA-1 | -4.43 | 2.37E-06 | NA | None | |
| HPOL_0002054401-mRNA-1 | -4.28 | 2.37E-06 | NA | None | |
| HPOL_0001537401-mRNA-1 | -4.25 | 7.51E-29 | NA | None | |
| HPOL_0001948701-mRNA-1 | -4.20 | 2.15E-4 | IPR019402:Frag1/DRAM/Sfk1 | 9 | Clade 4, 5 |
| HPOL_0002362201-mRNA-1 | -4.13 | 2.6E-2 | NA | 13 | Clade 5 |
| HPOL_0001938201-mRNA-1 | -4.04 | 1.51E-14 | NA | 11 | Clade 5 |
